# Neonatal hyperoxia induces sex-dependent pulmonary cellular and transcriptomic changes in an experimental mouse model of bronchopulmonary dysplasia

**DOI:** 10.1101/2022.07.12.499826

**Authors:** Sheng Xia, Lisandra Vila Ellis, Konner Winkley, Heather Menden, Sherry M. Mabry, Daniel Louiselle, Margaret Gibson, Elin Grundberg, Jichao Chen, Venkatesh Sampath

## Abstract

Hyperoxia (HOX) disrupts lung development in mice and causes bronchopulmonary dysplasia (BPD) in neonates. To investigate sex-dependent molecular and cellular programming involved in HOX, we surveyed the mouse lung using single cell RNA sequencing (scRNA-seq), and validated our findings in human neonatal lung cells *in vitro*. HOX-induced inflammation in alveolar type (AT) 2 cells gave rise to damage associated transient progenitors (DATP). It also induced a new subpopulation of AT1 cells with reduced expression of growth factors normally secreted by AT1 cells, but increased mitochondrial gene expression. Female alveolar epithelial cells had less EMT and pulmonary fibrosis signaling in HOX. In the endothelium, expansion of Car4+ EC (Cap2) was seen in HOX along with an emergent subpopulation of Cap2 with repressed VEGF signaling. This regenerative response was increased in females exposed to HOX. Mesenchymal cells had inflammatory signatures in HOX, with a new distal interstitial fibroblast subcluster characterized by repressed lipid biosynthesis and a transcriptomic signature resembling myofibroblasts. HOX-induced gene expression signatures in human neonatal fibroblasts and alveolar epithelial cells *in vitro* resembled mouse scRNA-seq data. These findings suggest that neonatal exposure to HOX programs distinct sex-specific stem cell progenitor and cellular reparative responses that underpin lung remodeling in BPD.

## INTRODUCTION

Mammalian lung development progresses sequentially through the embryonic, pseudoglandular, canalicular, saccular and alveolar stages ^1^. Establishment of the terminal gas exchange unit, the alveolus, lined by alveolar epithelial type 1 cells (AT1) and juxtaposed with capillary endothelial cells (EC) is the *summum bonum* of lung development. Lung ontogeny is temporally and spatially regulated by cell autonomous and non-autonomous signaling networks that specify cellular phenotypes, diversity, maturation and cell-cell interactions that establish the alveolar capillary barrier. In recent years, lung single cell RNA-sequencing (scRNA-seq) has revolutionized our understanding of the complexity and uniqueness of developmental programming of the cells that specify the alveolar niche ^2–7^. This fundamental knowledge is rapidly allowing us to interrogate deviant programming of cellular phenotypes underlying lung diseases using experimental animal models and human pathology samples.

Bronchopulmonary dysplasia (BPD) is a chronic lung disease that affects newborns born prematurely with canalicular or saccular lungs, not primed for gas exchange^8^. Preterm infants are exposed to high concentrations of oxygen (hyperoxia), that causes lung injury and disrupts lung development ^8,9^. Pathologically, a reduction in the alveoli with a pruning of vascular arborization, along with fibrosis and dysplastic lung development are well described. Signaling cascades related to the VEGF, TGFB, TP53, RhoA, IL1 and fibrosis pathways underlie the pathological changes observed in BPD^10–15^. While disturbances in these signaling pathways are ascribed to arise from alveolar epithelial, endothelial, mesenchymal and immune populations, transcriptomic networks that program cell-type specific alterations in phenotype specification, development and function in BPD are incompletely understood. We undertook a single cell survey in a hyperoxia model of experimental BPD to address these key questions. Strikingly, female preterm neonates are less susceptible to BPD in comparison with gestational age matched male preterm infants^16,17^. We also explored the mechanisms associated with decreased hyperoxia-induced lung injury in female neonatal mice at single cell resolution^18–21^.

ScRNA-seq studies have defined >35 lung cell types that can be grouped into 4 major cell lineages: the epithelium, endothelium, stromal cells and immune cells ^2,22^. The alveolar epithelium is comprised of cuboidal alveolar epithelial type 2 (AT2) cells that secrete surfactant, and serve as progenitor cells in the adult lung ^23,24^. AT1 cells are thin, flat cells that cover the alveolar surface, and facilitate gas exchange and alveolarization^25,26^. In models of adult lung injury, AT2 cells serving as progenitor cells repopulate the AT1 population^5,24,27^. Injury-induced AT2 inflammation represses AT2 differentiation to AT1 and leads to alveolar epithelial-to-mesenchymal transition (EMT) ^28,29^. TP53 and TGFB inhibit AT2-AT1 intermediate cell maturation to AT1 cells, favoring lung fibrosis^23,29^. Whether hyperoxia (HOX)-induced lung injury programs changes in AT2 to restore the AT1 population in BPD is unknown. Lung capillary EC specify into capillary endothelial cell type 2 (Cap2, or aCap) that lie opposed to AT1, participate in gas exchange, and capillary endothelial cell type 1 (Cap1, or gCap), which have distinct vasomotor, immune and regenerative function^3,30–33^. *Car4* expressing Cap2 EC are known to respond to paracrine signals, and might be key to EC and alveolar regeneration ^31^. However, since Cap2 do not proliferate, it is accepted that Cap1, which retain proliferative and progenitor properties, give rise to Cap2 during lung injury. How HOX in the neonatal lung programs cellular and transcriptional networks important for EC recovery is not known. Alveolar myofibroblasts (AMF) (myofibroblast 1), ductal myofibroblasts (DMF) (myofibroblast 2), distal interstitial fibroblasts (DIF) (lipofibroblasts) and proximal interstitial fibroblasts (PIF) (matrix fibroblasts) are four major fibroblasts involved in alveolarization and alveolar maintenance^3,34–36^. Excessive proliferation of injury-transformed fibroblasts and extracellular matrix accumulation leads to lung fibrosis^37^. The impact of HOX on fibroblasts subpopulations and transcriptional networks that contribute to matrix remodeling in human and experimental BPD was elucidated herein.

In this study, we investigated the cell-type specific alterations in transcriptomic networks that program injury, repair and deviant development in BPD using a model of experimental BPD in which mice are exposed to hyperoxia (85%) from P1-P14. Combining scRNA-seq data with validation studies in mice and human lung cell lines we show that experimental BPD programs unique phenotypic changes in lung alveolar epithelial cell, capillary EC, fibroblast, and immune cell populations that underpin injury/adaptive responses in HOX. We also investigated sex-specific transcriptomic changes in the injury-responsive cells that might explain the decreased vulnerability or augmented regenerative response to hyperoxia injury seen in female infants.

## RESULTS

### Lung cellular composition of the developing mouse lung at P14 after room air and hyperoxia exposures revealed by scRNA-seq

We and others have shown that HOX induces alveolar simplification with arrested vascular and alveolar development ^38–40^. To study the effect of HOX on lung cellular lineages and subpopulations, we used a well-established model of experimental BPD. We exposed newborn C57BL/6 mice to room air (RA; 3 males and 3 females) and hyperoxia (HOX; 3 males and 3 females) from P1 to P14 (Fig 1A). We used western blotting, immunofluorescence studies and qRT-PCR in independent mice (4 males and 4 females) and human neonatal lung cells/lines to validate our scRNA-seq findings (Fig 1A). HOX reduced radial alveolar counts and increased mean linear intercepts by 40% (Sup. Fig. 1A and 1B). To examine cell type-specific changes in the transcriptome associated with hyperoxia, we performed scRNA-seq (Fig. 1B) and identified four major cell lineages: *Nkx2-1*^+^ epithelial cells, *Cdh5*^+^ endothelial cells, *Col3a1*^+^ stromal cells and *Ptprc*^+^ immune cells^3,30^ (Fig. 1C). We identified marker genes for AT1 cells (*Aqp5*), AT2 cells (*Sftpc*), ciliated epithelial cells (*Foxj1*) and club cells (*Scgb1a1*) in the epithelial cell cluster. We identified lineage markers for Cap1 EC (*Plvap*), Cap2 EC (*Car4*), big blood vessel EC (*Vwf*) and lymphatics marker (*Prox1*) in EC cluster. Markers for stromal cells associated with the three axes–vascular, epithelial and interstitial were identified, which encompass the pulmonary smooth muscle cells, pericytes and fibroblast subpopulations ^36^. Finally, markers for distinct immune cell populations were identified (Fig. 1D). Because we identified new subclusters within major cell clusters in HOX, a detailed description of subpopulations is described within their respective sections.

**Figure 1.**
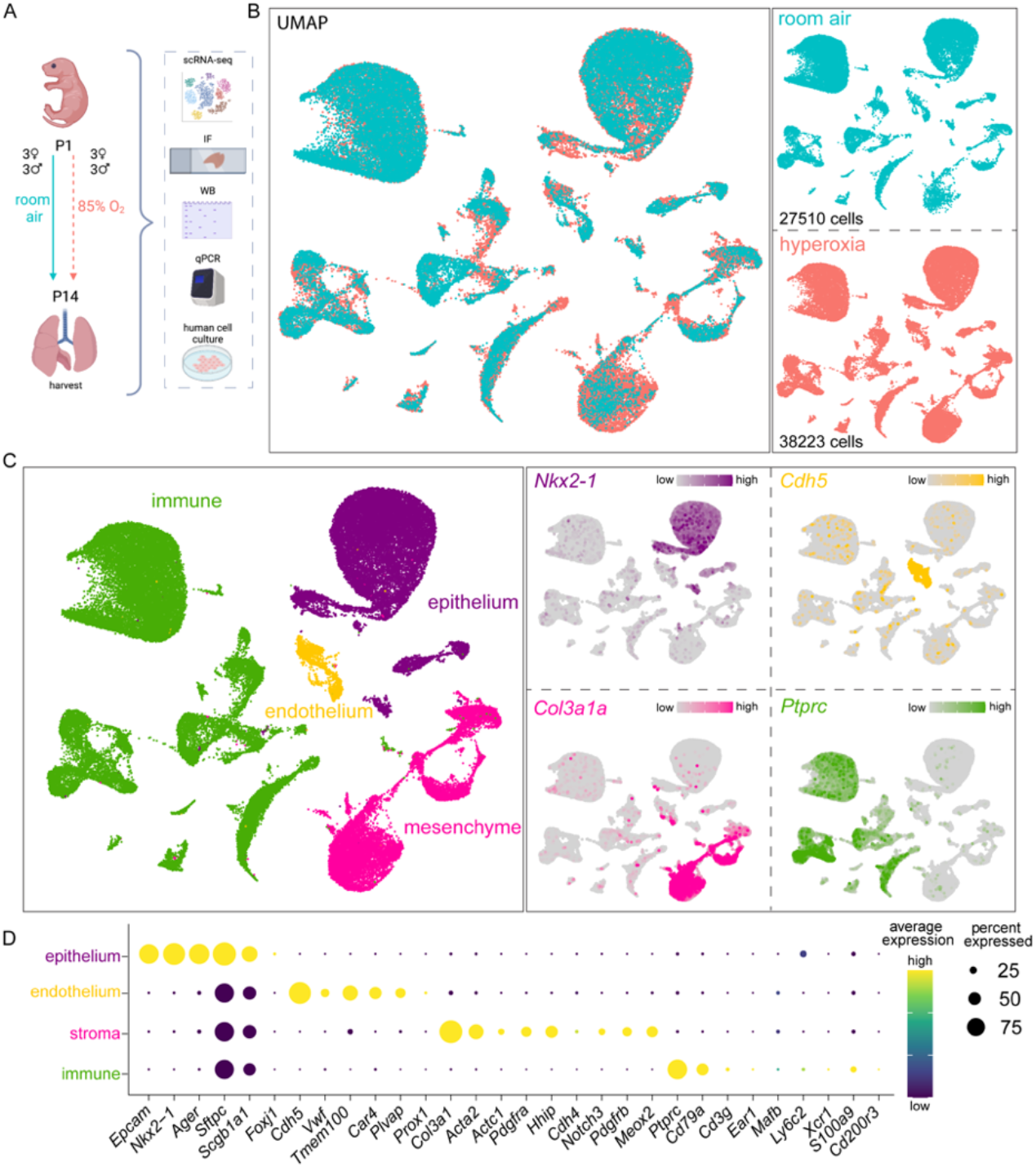
Lung cellular architecture in room air and hyperoxia at P14 by scRNA-seq. A) Illustration generated with permission on BioRender of experimental design: mouse pups were exposed to 85% oxygen or RA from P1 to P14, and the lungs were used for single cell RNA-seq (scRNA-seq), immunofluorescence (IF), whole lung lysate qRT-PCR and western blotting. Human primary cells were used for validation of mouse scRNA-seq data. B) Left: scRNA-seq UMAP with merged treatment conditions, RA (blue) and HOX (pink), showing overlapping clusters. Right: UMAPs of cells in each condition show similarities and differences, with cell numbers specified in each case. 3 females and 3 males were sequenced per group. C) Seurat unbiased clustering grouped all cells into 37 clusters which we attributed to four cell lineages shown in UMAP (left). Epithelial, endothelial, stromal, and immune cell identities were determined by the expression of *Nkx2-1*, *Chd5*, *Col3a1*, and *Ptprc*, respectively, shown in feature plots (right). D) Dot plot depicting expression level (dot color) and percentage of expression (dot size) of markers used to identify subpopulations within each major cell lineage.

### Alterations of canonical pathways, upstream genes and functions indicate upregulation of oxidative phosphorylation, NRF2, TP53 and TGFβ1 signaling in major cell lineages

BPD is associated with several changes in developmental, cell injury, fibrosis, and inflammatory pathways^8,9^. We therefore analyzed our scRNA-seq data using Ingenuity Pathway Analysis (IPA) (Qiagen) for signaling and functional pathways that were significantly altered in HOX compared to room air (RA), and were shared across major cell lineages (Fig. 2A-C). Oxidative phosphorylation and NRF2-mediated oxidative stress response were upregulated in all cell lineages (Fig. 2A). Fibrosis signaling was upregulated in all cell lineages except for stromal cells. NRF2-mediated oxidative stress response activates expression of antioxidant response elements (ARE) that protect the lung from oxidative stress caused by HOX ^41,42^. TGFβ signaling is important for lung development, but is also a key mediator of fibrosis ^43^, was altered in all lineages (Fig. 2B). TP53, a regulator of lung injury and fibrosis ^13,44^, was also upregulated (Fig. 2B). These changes support the increased fibrosis signaling found in HOX (Fig. 2A). Integrin signaling and actin nucleation by ARP-WASP complex were upregulated in epithelial cells and immune cells, which is consistent with increased cell migration in function analysis (Fig. 2A, 2C).

**Figure 2.**
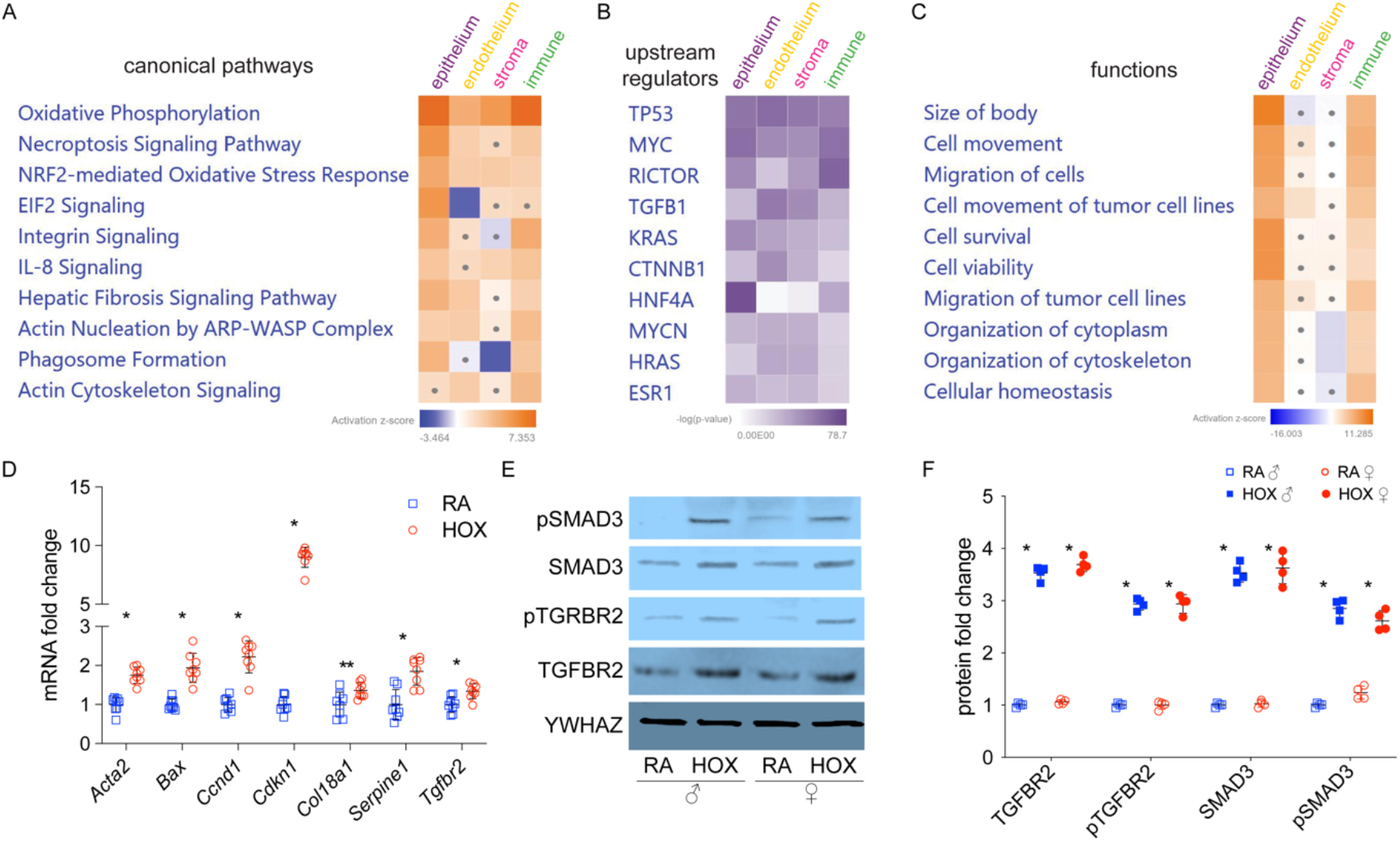
Alterations of canonical pathways, upstream regulators, and functions in hyperoxia. scRNA-seq differential gene expression data was used to generate IPA heatmaps of top 10 canonical pathways (A), upstream regulators (B) and functions (C) altered by HOX. Dots in heatmaps represent values that had a |Z-score| of <2 (non-significant). D) Mouse whole lung qRT-PCR shows fibrosis marker genes (*Ccnd1*, *Acta2*, *Serpine1*, and *Col18a1*), TGFB1 receptor (*Tgfbr2*) and its downstream genes (*Acta2*, *Tgfbr2*, *Serpine1*, and *Cdkn1a*) are upregulated by HOX treatment. n=8 per group; *p<0.005; **p<0.05 E) Mouse whole lung immunoblotting shows expression and phosphorylation of TGFRB2 and SMAD3 are stimulated by HOX in males and females, with densitometry shown graphically F), n=4 per group; *p<0.05.

In independent experiments, we collected whole lung samples from neonatal littermates exposed to HOX or RA from P1 to P14 (n=8/group, 4 in each sex). We validated genes associated with signaling of TGFβ and fibrosis. Quantitative reverse transcription PCR (qRT-qPCR) showed that fibrosis marker genes (*Ccnd1*, *Acta2*, *Serpine1* and *Col18a1*), TGFB1 downstream genes (*Acta2*, *Tgfbr2*, *Serpine1* and *Cdkn1a*) and TP53 signaling genes (*Serpine*, *Bax*, *Cdkn1a*) in whole mouse lungs with HOX treatment (Fig. 2D) ^45–52^. Because our gene signatures suggested activation of TGFβ and fibrosis signaling, we performed western blotting (WB) on whole lung lysates to demonstrate that expression and phosphorylation of TGFBR2 and SMAD3, canonical markers of TGFβ activation, were induced in HOX (Fig. 2E and 2F). YWHAZ was used for normalization as β-actin expression changed with HOX. These data show that HOX induces TGFβ and pulmonary fibrosis signaling among several lung cell types, consistent with whole lung data reported before ^12,13,43,44^.

### Hyperoxia induces damage-associated transient progenitors (DATP) and immature AT1 (iAT1) populations

We next analyzed the major cell lineages individually, starting with epithelial cells. In *Nkx2-1*^+^/*Epcam*^+^/*Cdh1*^+^ epithelial cells, we identified 3 main clusters: *Etv5*^-^/*Pdpn*^+^ AT1 cells, *Etv5*^+^/*Sftpc*^+^ AT2 cells, and *Etv5*^+^/*Pdpn*^+^/*Sftpc*^+^ AT2/AT1 cells. We also identified minor populations of *Foxj1*^+^ bronchiolar ciliated cells and *Scgb1a1*^+^ basal cells (data not shown), but we focused on alveolar epithelial cells. Based on gene expression analysis, we categorized AT2 cells into 4 subclusters in RA: *Lyz1*^+^AT2, AT2, *Mki67*^+^ cycling AT2 (cAT2), and *mt-Nd1/2/4^high^* mitochondrial AT2 (mAT2) (Fig. 3A). cAT2 cells express proliferation-related genes, such as *Mki67* and *Cdk1* (Fig. 3A) ^28^. *Lyz1*^+^ AT2 cells are enriched with the lysozyme gene *Lyz1* (Fig. 3A), and have been previously described^7^. Comparing *Lyz1*^+^ AT2 to AT2 cells, IPA upstream analysis revealed that ETV5 signaling, which controls *Lyz1* expression in lung AT2 cells^53^, was upregulated in *Lyz1*^+^ AT2 cells (Sup. Fig. 2A). IPA pathway analysis also identified downregulated mTOR, actin cytoskeleton and pulmonary healing signaling pathways, as well as upregulated Hippo signaling in *Lyz1*^+^ AT2 (Sup. Fig. 2B), suggesting less proliferation, migration, and alveologenesis potential. The mAT2 population, which has not been described before, express reduced surfactant genes (*Sftpa*, *Sftpb* and *Sftpc*) and less AT2 linage marker genes (*Etv5*, *Abca3* and *Cebpa*)^28,53,54^, but more mitochondrial genes, such as *mt-Nd1/2/4* (Fig. 3A). IPA upstream and canonical pathways analysis demonstrated less ETV5 signaling (Sup. Fig. 2C) and more mitochondria biogenesis - part of upregulated Sirtuin signaling - in mAT2 compared to AT2 cells (Sup. Fig. 2D).

**Figure 3.**
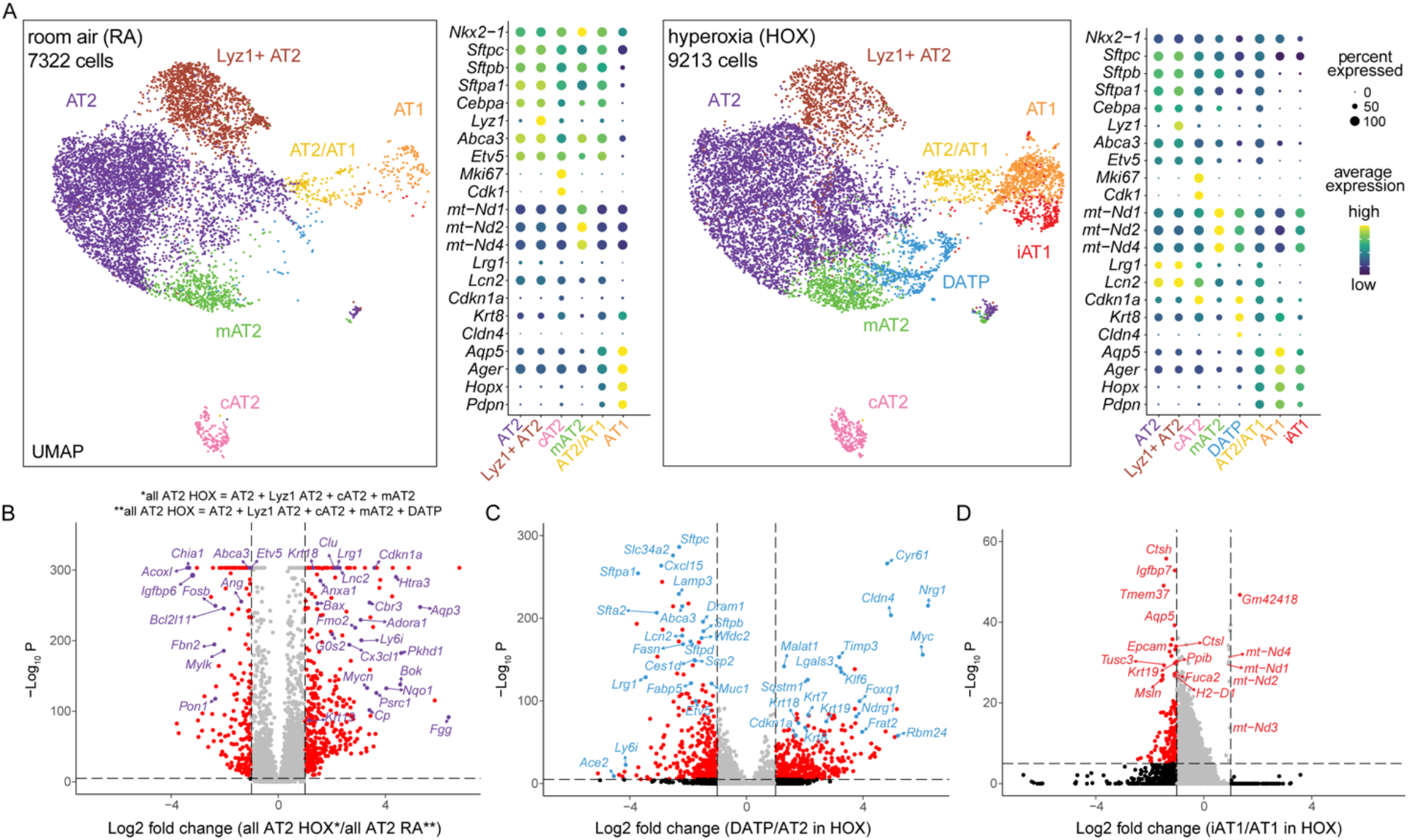
scRNA-seq revealed alveolar epithelial diversity and transcriptomic changes in hyperoxia. A) UMAPs showing the distinct alveolar epithelial clusters identified, along with dot plots featuring the marker genes used to characterize them. In RA (right), we found AT2 cells consisted of four subpopulations (AT2, Lyz1^+^ AT2, mAT2 and cAT2), and found two other clusters including AT2/AT1 and AT1 cells. In HOX (left), two additional subpopulations emerged, a cluster of AT2 cells labeled DATP (blue), and a cluster of AT1 cells labeled iAT1 (red). Volcano plots comparing gene expression between (B) all AT2 cells in HOX vs AT2 cells in RA, (C) DATP and AT2 in HOX, (D) iAT1 and AT1 in HOX. Down/upregulated genes are shown in red, while some genes of biological relevance have been highlighted.

Interestingly, we found major changes in the AT2 transcriptome in HOX (Fig. 3A). The majority of AT2 clusters, including AT2 and *Lyz1*^+^AT2, were enriched for inflammation-related genes (*Lcn2* and *Lrg1*), but expressed less AT2 lineage markers (*Etv5*, *Abca3*, and *Cebpa*) (Fig. 3A). A distinct *Sftpc^low^ /Krt8*^+^/*Cldn4*^+^ AT2 subcluster emerged in HOX that was not evident in normal developing lungs (Fig. 3A). These cells have been labelled as damage-associated transient progenitors (DATP) in a mouse model of bleomycin-induced fibrosis ^28^. A comparison between all AT2 cells present in HOX vs RA, showed less AT2 lineage marker genes (*Etv5* and *Abca3*), more AT2 to AT1 differentiation marker genes (*Krt18* and *Krt19*) and more inflammation-related genes (*Lcn2* and *Lrg1*) ^28,53–56^ (Fig. 3B). In turn, DATP cluster showed less AT2 lineage markers (*Etv5*, *Abca3*, *Sftpb* and *Sftpc*) and more AT2 to AT1 differentiation marker genes (*Krt7*, *Krt8*, *Krt18* and *Krt19*) when compared to AT2 in HOX ^28^,^53–57^.

AT2/AT1 cells express AT2 lineage maker genes (*Etv5*, *Abca3* and *Cebpa*), and at a lower level several AT1 cell marker genes (*Aqp5*, *Ager*, *Hopx* and *Pdpn*) in RA (Fig. 3A). The AT2/AT1 population had similar signatures in RA vs HOX. We identified one AT1 cell cluster that was enriched with classical AT1 marker genes (*Hopx*, *Pdpn* and *Aqp5*), and did not express AT2 marker genes (*Sftpc*, *Etv5* and *Abca3*) in RA or HOX. Interestingly, one new subcluster of AT1 cells, which we labelled immature AT1 (iAT1), emerged in HOX (Fig. 3A). Notably, iAT1 had increased expression of mt-Nd1/2/3/4 but less AT1 mature marker genes (*Aqp5*, *Epcam*) when compared to AT1 in HOX (Fig. 3D). IPA analysis showed less senescence and lung fibrosis signaling in iAT1 cells compared to AT1 in HOX (Sup. Fig. 2E). iAT1 cells also express less *Vegfa*, *Pdgfa*, *Fgf18* and *Fgf1* when compared to AT1 in HOX (Sup. Fig. 2F). *Vegfa* and *Pdgfa*, play critical roles in alveolarization ^58–61^, while *FGF18* regulates key developmental events during pulmonary alveolarization, and *FGF1* stimulates epithelial proliferation^62,63^. These transcriptional changes in iAT1 cells suggest a decreased ability to contribute to alveolarization.

To confirm these interesting findings, we performed immunofluorescence (IF) on lung sections from HOX and RA treated pups. We confirmed the general inflamed signature of AT2 cells by demonstrating that LCN2 was increased in AT2 in HOX and co-localized with SPC staining (Fig. 4A). KRT8^high^/SPC^+^ staining showed DATP emerged in HOX (Fig. 4B), but was not present in RA. SPC/PDPN staining revealed three clusters of alveolar epithelial cells, which are SFTPC^+^ AT2 cells, PDPN^+^ AT1 cells and PDPN^+^/SFTPC^+^ AT2/AT1 cells, the latter were increased in HOX (Fig. 4C). iAT1 cells were visualized with MT-ND1/PDPN staining in HOX (Fig. 4D).

**Figure 4.**
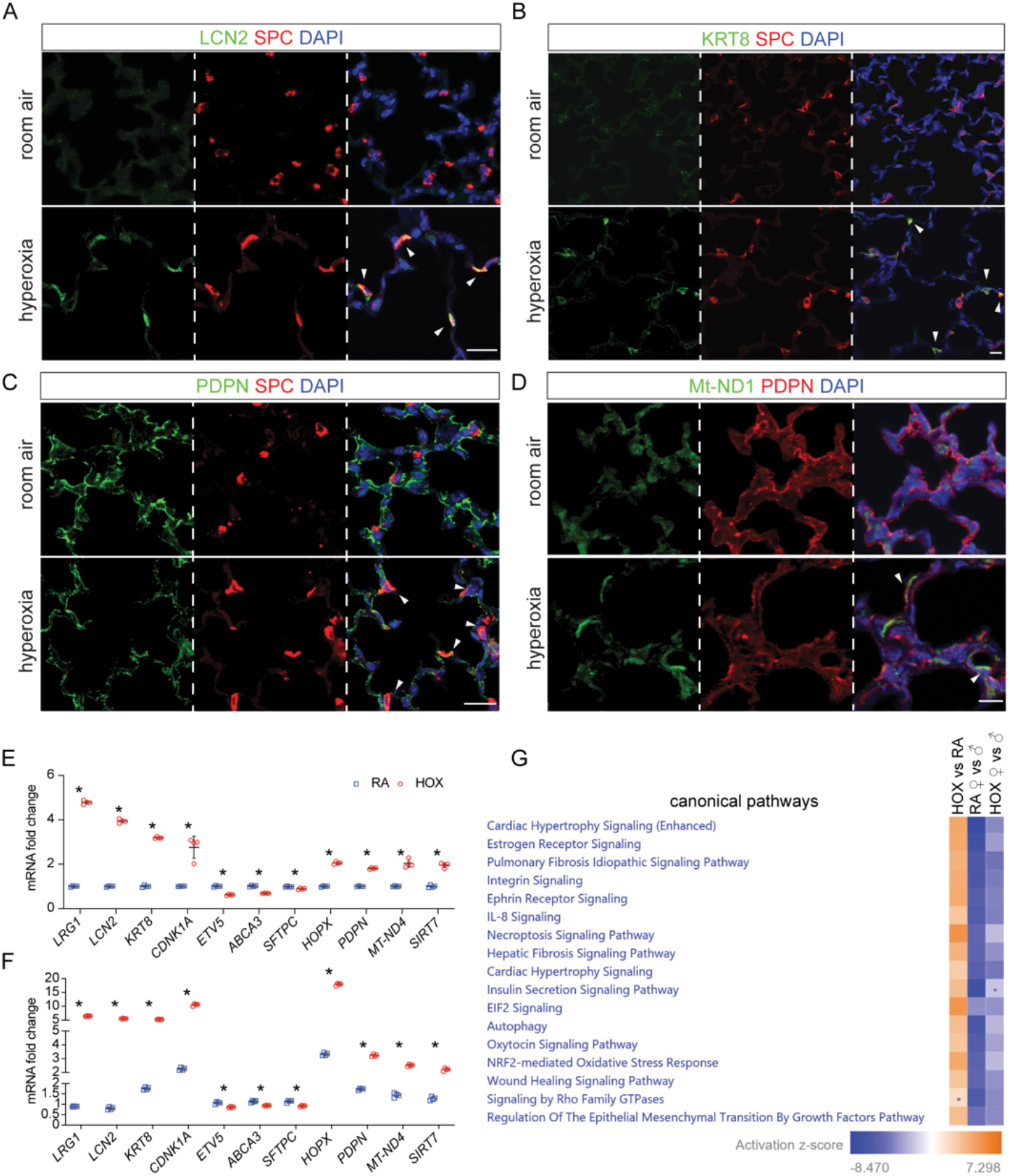
Mouse *in vivo* immunostaining and human cell culture confirmed alveolar epithelial diversity and alterations in hyperoxia. (A-D) Immunofluorescence of mouse lung sections following RA or HOX treatment from P1-P14; scale bar: 10μm. A) AT2 cells, labeled with SPC, show co-expression of inflammation marker LCN2 (arrowhead) in HOX. B) Emergence of DATP cells is seen in the alveolar region after HOX, where some AT2 cells which express SPC, also express KRT8 (arrowhead). C) PDPN and SPC co-localization (arrowhead) demonstrates increased AT2/AT1 cells in HOX. D) HOX-induced iAT1 cells can be seen by co-expression of Mt-ND1 and AT1 marker PDPN (arrowhead). E) RNA obtained from human pulmonary alveolar epithelial cells (HPAEpiC) exposed to 48hr of RA or HOX (85% O_2_) was used to quantify *LRG1*, *LCN2*, *KRT8*, *CDNK1A*, *ETV5*, *ABCA3*, *SFTPC*, *HOPX*, *PDPN*, *mt-ND4*, and *SIRT7* with qRT-PCR. n=4 per group; *p<0.05. F) RNA obtained from 48hr RA and HOX treated HPAEpiC, which were co-cultured with immortalized human pulmonary fibroblasts (HPF-Im), was used to quantify *LRG1*, *LCN2*, *KRT8*, *CDNK1A*, *ETV5*, *ABCA3*, *SFTPC*, *HOPX*, *PDPN*, *mt-ND4*, and *SIRT7* with qRT-PCR. n=4 per group; *p<0.05. G) IPA canonical pathway analysis heatmap showing several upregulated pathways in all alveolar epithelial cells in HOX; pulmonary fibrosis and wound healing signaling pathways were upregulated, but less so in females than in males. Dots in heatmaps represent values that had a |Z-score| of <2 (non-significant).

IL1β and HIF1α have been shown to induce AT2 differentiation to AT1 via DATP, while TGFβ signaling represses AT1 maturation ^28,55^. IPA upstream analysis showed that IL1 signaling is activated in Lyz1+ AT2, AT2, mAT2 and AT2/AT1 cells, while HIF1α signaling and TGFβ signaling was upregulated in mAT2 and AT2/AT1 cells in HOX (Sup. Fig. 2G). Upregulated TP53 signaling and HIF1α signaling in DATP was revealed by comparing to AT2 in HOX (Sup. Fig. 2H). Transcriptomic signatures of IL1R, HIF1α, and TGFβ activation seen in AT2 subpopulations with neonatal HOX resemble those described in adult mouse models of bleomycin-induced fibrosis ^28^, involved in AT2 reparative responses. These data show that HOX programs AT2 to a transitional “stem-cell like fate” described in adult models of fibrosis^64^.

To validate our mouse data, we exposed neonatal human primary alveolar epithelial cells (HPAEpiC) isolated from the neonatal lung (ScienCell) to HOX or normoxia for 48 hours, and then examined marker gene expression of AT2 and AT1 subsets identified in HOX using qRT-PCR. We found that markers of inflammation (*LRG1* and *LCN2*), DATP signature (*KRT8*), mitochondria biogenesis signature (*mt-ND4* and *SIRT7*^65^) were upregulated in response to HOX (Fig. 4E). cAT2 and DATP shared marker gene *CDNK1A*, was also upregulated in HOX (Fig. 4E). AT2 marker genes (*ETV5*, *ABCA3* and *SFTPC*) were slightly downregulated and AT1 marker genes (*HOPX* and *PDPN*) were upregulated (Fig. 4E). Interestingly, when HPAEpiC were co-cultured with human neonatal lung fibroblasts most of these changes were accentuated (Fig. 4F). These data indicated that HOX induces signatures of mitochondrial biogenesis and DATP cell autonomously, laying the foundation for differentiation of AT2 to AT1.

IPA analysis showed that pulmonary fibrosis and epithelial to mesenchymal transition (EMT) signaling were upregulated in AT cells in HOX (Fig. 4G). We explored female vs male differences in our scRNA-seq data and found that there was less pulmonary fibrosis and EMT signaling in females, in both RA and HOX (Fig. 4G). The decreased activation of lung fibrosis and EMT could suggest more programming of AT2 to AT1 rather than fibrosis in female mice exposed to HOX.

### New subpopulation of Car4+ Cap2 endothelial cells emerges in hyperoxia

We analyzed EC in RA and identified six previously known clusters: *Cdh5*^+^/*Plvap*^+^/*Aplnr*^+^/*Vwf*^-^Cap1, *Cdh5*^+^ /*Car4*^+^/*Apln*^+^/*Plvap*^-^ Cap2, *Cdh5*^+^/*Vwf*^+^/*Nr2f2*^+^ venous EC, *Cdh5*^+^/*VWF*^+^/*Gja5*^+^ arterial EC, *Cdh5*^+^/*Prox1*^+^ lymphatic EC and *Cdh5*^+^/*Mki67*^+^ cycling EC (cEC) (Fig. 5A) ^3,30–33^. The proportion of capillary EC identified as Cap1 decreased in HOX, while the Cap2 proportion increased (Fig. 5A). Dot plots showed that *Car4* and *Apln* expression was upregulated, while *Aplnr* and *Plvap* were downregulated in Cap1 in HOX (Fig. 5A), which may indicate a skewing of capillary transcriptomic signature from Cap1 to Cap2 in HOX. Interestingly, in adult mouse models of influenza-induced lung injury, Cap1 cells are considered to be progenitor cells, giving rise to Cap2 (CAR4^+^) cells after infection-induced injury^33^. This is also known to be the case during normal lung ontogeny ^31,32^. Unlike in RA, the Cap2 population in HOX appeared heterogenous with an immature Cap2 (iCap2) subcluster (Fig. 5A). These cells were specifically enriched for *Lilr4b*, *Ryr2* and *Gpx3* (Fig. 5A), had less *Car4* expression but more *Apln* expression compared to regular Cap2, and did not express the Cap1 markers *Plvap* and *Aplnr* (Fig. 5A). Interestingly, the comparison between iCap2 and Cap2 in HOX showed significant downregulation of *Icam2*, a marker of lumenized vessels, further indicating these cells are immature and may be undergoing vascular remodeling (Sup. Fig 3A) ^66^. IPA analysis revealed that VEGF signaling, which is important for Cap2 differentiation ^31^, was repressed in iCap2 compared to Cap2 (Sup. Fig. 3B).

**Figure 5.**
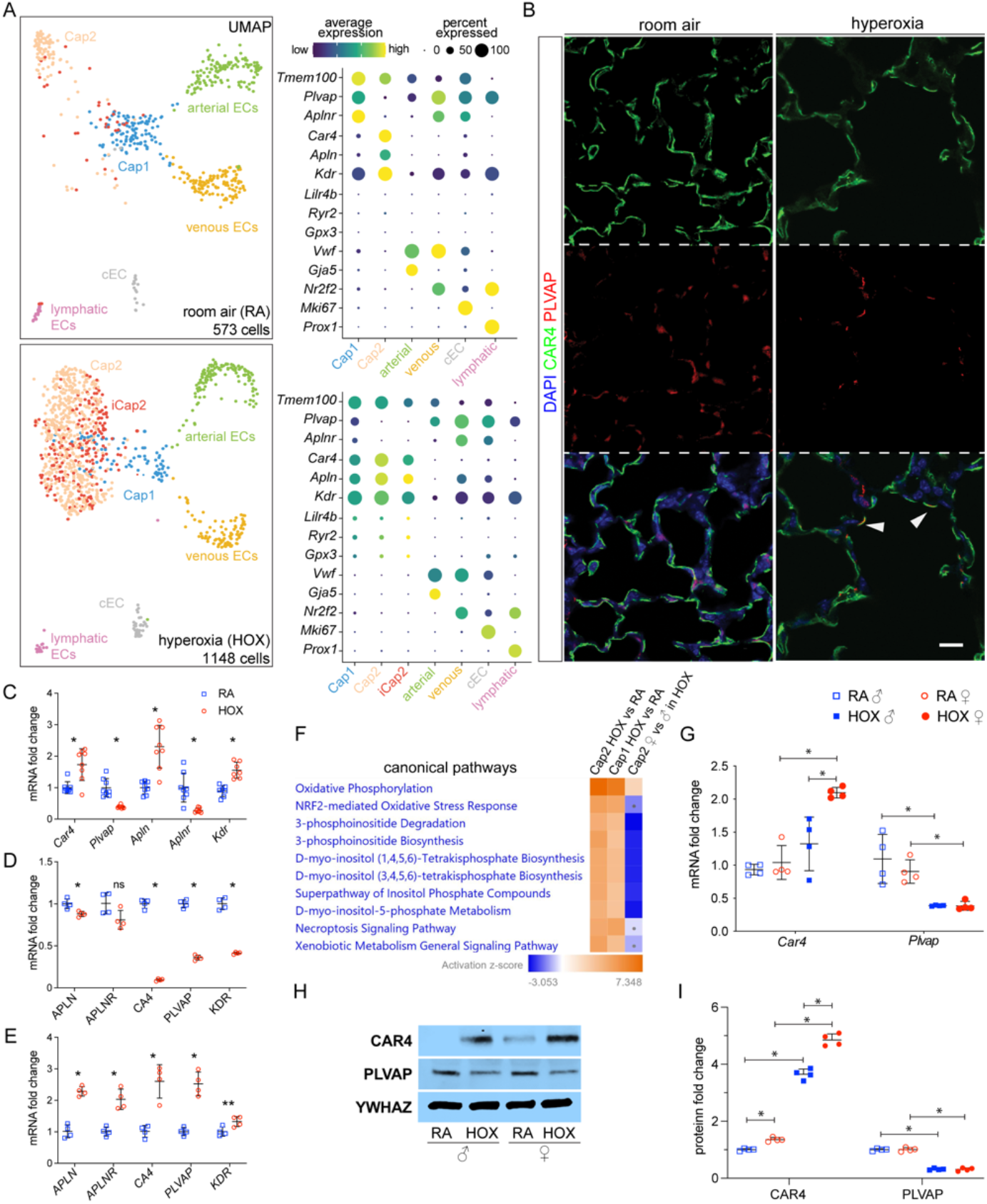
Hyperoxia induces a new subpopulation of *Car4*^+^ Cap2 cells and causes sex-specific transcriptomic changes in the endothelium. A) UMAPs of endothelial cell (EC) clusters in RA (top) and HOX (bottom). In RA, six clusters were identified using the marker genes shown in dot plots: *Plvap*^+^ Cap1, *Car4*^+^ Cap2, *Gja5*^+^ arterial EC, *Nr2f2*^+^ venous EC, *Mki67*^+^ cEC, and *Prox1*+ lymphatic EC. In HOX, an additional subpopulation of Cap2 emerged which we named iCap2 (red). B) Immunostaining of mouse lung sections showing a general reduction in vessel density in HOX compared to HOX. Co-expression of CAR4 and PLVAP was also observed in HOX (arrowhead). Scale bar: 10μm. C) Mouse whole lung qRT-PCR depicts upregulated expression of Cap2 EC marker genes (*Car4*, *Apln*, and *Kdr*) and downregulated expression of Cap1 EC marker genes (*Plvap* and *Aplnr*) in HOX compared to RA. n=8 per group; *p<0.005. D) RNA obtained from immortalized human pulmonary microvascular endothelial cells (HPMEC-Im) exposed to 7 days of RA or HOX (85% O_2_) was used to quantify *APLN*, *APLNR*, *CA4*, *PLVAP*, and *KDR* with qRT-PCR. n=4 per group; *p<0.05. E) RNA obtained from HPMEC-Im exposed to 7 days of RA or HOX (85% O_2_) that were co-cultured with immortalized human pulmonary fibroblasts (HPF-IM), was used to quantify *APLN*, *APLNR*, *CA4*, *PLVAP*, and *KDR* with qRT-PCR. n=4 per group; p<0.005; **p<0.05. F) Heatmap of top 10 HOX-altered canonical pathways analyzed with IPA showing that female Cap2 EC had increased oxidative phosphorylation compared to their male counterpart, while other pathways were downregulated. The dots in heatmaps represent values that had a |Z-score| of <2 (non-significant). G) Mouse whole lung qRT-PCR showing decreased *Plvap* mRNA expression in HOX samples in both females and males, but higher *Car4* mRNA expression in females only. n= 4 per group; *p<0.05. H) Mouse whole lung immunoblotting reveals that HOX represses PLVAP expression and stimulates CAR4 expression in both females and males, but stimulates more CAR4 expression in females vs males, with densitometry shown graphically in I), n= 4 per groups; *p<0.05.

Next, we proceeded to validate our scRNA-seq findings in independent RA and HOX-treated mouse samples with IF. CAR4^+^ EC (Cap2) cover most of alveolar surface abutting the air interface, while PLVAP+ EC (Cap1) localize to the opposite side of the alveolar capillary interface ^31,32^ (Fig. 5B). In addition to a general reduction in vessel density, we observed more co-localization between CAR4 and PLVAP staining in HOX (Fig. 5B), which would suggest a greater proportion of Cap1 cells may be transitioning to Cap2 in HOX. *Car4, Apln* and *Kdr* were enriched in Cap2, while *Plvap* and *Aplnr* were enriched in Cap1, as expected (Fig. 5A). Whole lung RNA expression by qRT-PCR demonstrated increased expression of *Car4*, *Apln* and *Kdr*, but decreased expression of *Aplnr* and *Plvap* (Fig. 5C) in HOX, which was also confirmed by our immunoblotting data (Fig. 5H and 5I) showing increased CAR4 and decreased PLVAP in HOX. These results suggest an increase in the proportion of Cap2, and decreased Cap1 as noted in our scRNA-seq UMAP (Fig 5A).

To validate our mouse data in human lung EC, we used an immortalized fetal human pulmonary microvascular endothelial cells (HPMEC-Im) that we had generated before ^67^. We exposed HPMEC-Im to HOX (85%) or normoxia for 7 days in the presence or absence of immortalized fetal human pulmonary fibroblasts (HPF-Im) to test cell-autonomous changes in gene expression in HOX. Surprisingly, qRT-PCR demonstrated that HOX suppressed Cap2 and Cap1 markers [*APLN*, *APLNR*, *CA4* (orthologous to mouse *Car4*), *PLVAP* and *KDR*] in the absence of fibroblasts (Fig. 5D). However, when co-cultured with HPF-Im for seven days, expression of both Cap2 and Cap1 markers were increased in HOX (Fig. 5E). These data suggest fibroblasts are required for the induction or maintenance of *CAR4* expression in HOX.

We then did IPA analysis on gene expression profiles in Cap2 and Cap1 in HOX vs RA. The top differentially expressed canonical pathways in HOX included oxidative phosphorylation, NRF2-oxidative stress response genes, and inositol biosynthesis signaling (Fig. 5F). Comparing female vs male Cap2 population showed that female Cap2 cells had increased oxidative phosphorylation compared to male Cap2 cells, while other pathways were suppressed (Fig. 5F). We did not have enough Cap1 cells in HOX to do a male vs female comparison.

qRT-PCR showed that *Car4* mRNA expression was upregulated by HOX in females, but not in males, and its expression was higher in female than in males in HOX (Fig 5G). *Plvap* mRNA expression was downregulated by HOX in both males and females (Fig. 5G). Immunoblotting demonstrated that CAR4 protein levels were upregulated by HOX in both females and males, and its expression was higher in female than in males in HOX, while PLVAP protein expression was downregulated by HOX in both males and females (Fig 5H and 5I). This is suggestive of a more robust reparative EC response with increased transitioning of Cap1 to Cap2.

### Hyperoxia induces inflammation signature in stromal cells while specifying a fibrotic fibroblast subpopulation

In RA, we classified *Col3a1*^+^ stromal mesenchymal cells into 3 groups based on their proximity to vascular or epithelial structures nearby or “axes”, according to terminology published recently^36^. The vascular axis consists of *Notch3^high^/Gap43*^+^/*Pdgfrb^high-^* pericytes (PC) and *Notch3^high^/Gap43*^-^/*Acta2*^+^ vascular smooth muscle cells (VSMC); the epithelial axis is formed by *Pdgfra^high^/ Lgr6^-^/Hhip*^+^/*Cdh4*^+^ alveolar myofibroblasts (AMF), *Pdgfra*^+^/*Lgr6*^+^/*Hhip^high^/Cdh4^high^* ductal myofibrobltasts (DMF) and *Notch3*^+^/ Actc1^+^/*Lgr6*^+^ airway smooth muscle cells (ASMC); the interstitial axis includes *Pdgfra*^+^/*Twist2*^+^/*Il33*^+^/*Col1a1^high^*/ *Col1a2^high^* proximal interstitial fibroblasts (PIF) and *Pdgfra*^+^/*Wnt2*^+^/*Tcf-21^high^/G0s2^high^* distal interstitial fibroblasts (DIF) ^3,30,35,36^. (Fig. 6A).

**Figure 6.**
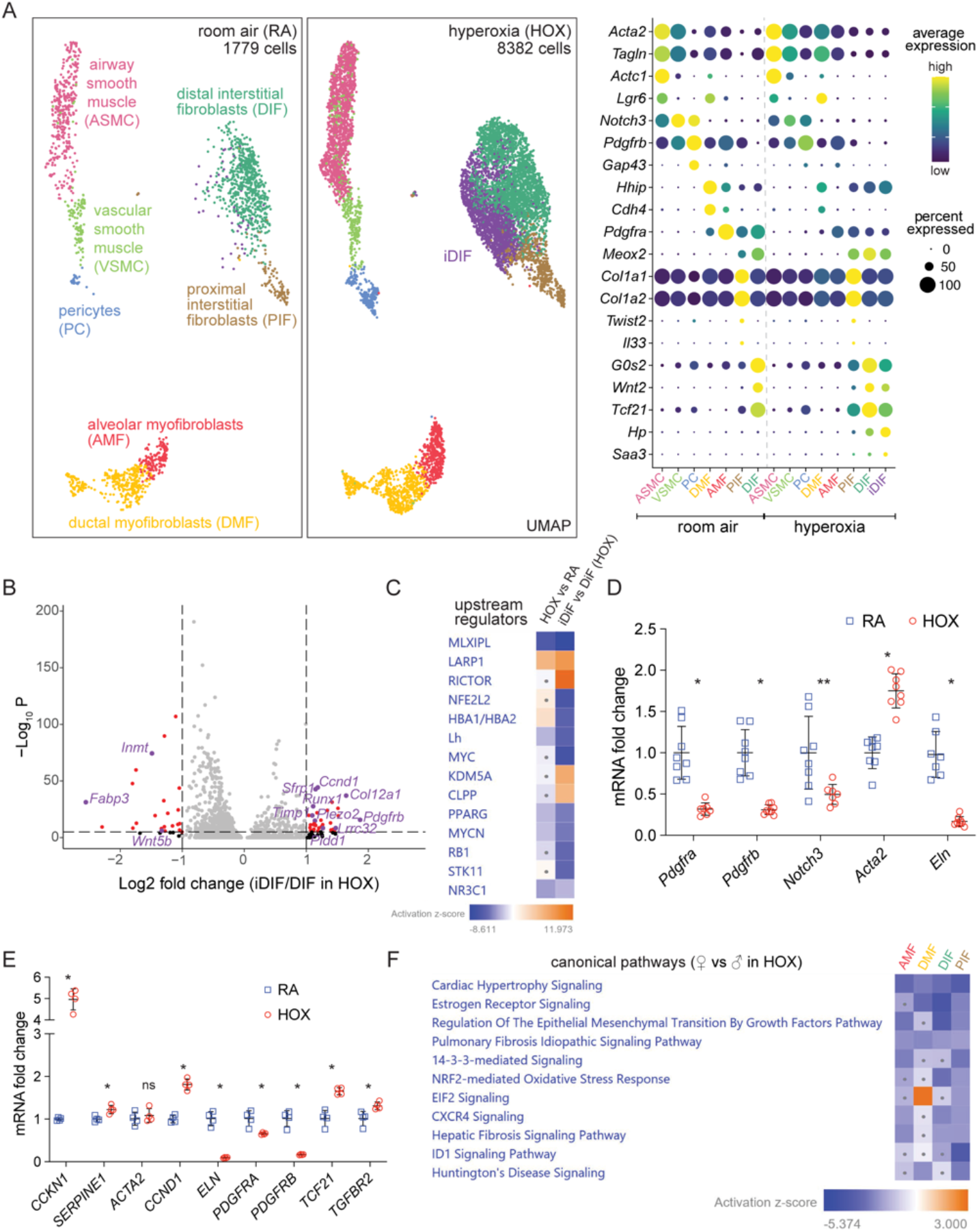
New population of distal interstitial fibroblasts emerges in hyperoxia and shows sex-specific characteristics. A) UMAP of stromal cell showing 7 clusters in RA (left) identified as DIF, PIF, AMF, DMF, PC, VSMC, and ASM. In HOX (middle), a new cluster emerged labeled as iDIF (purple). Dot plot showing marker genes used for identification of each population (right). B) Volcano plot shows gene expression differences between iDIF and DIF in HOX. Significantly affected genes are shown in red, with some biological relevant genes highlighted in purple. C) IPA analysis revealing downregulation of MLXIPL and PPARG signaling in HOX, while these two upstream regulators are even more repressed in iDIF vs DIF. The dots in heat maps represent values that had a |Z-score| of <2 (non-significant). D) Mouse whole lung qRT-PCR reveals downregulated expression of different stromal cluster marker genes, such as *Pdgfra*, *pdgfrb*, *Notch3*, and *Eln*, and upregulated Acta expression in HOX. n=8 per group; *p<0.001; **p<0.02. E) Immortalized human pulmonary fibroblasts (HPF-IM) in culture were exposed to 7 days of RA or HOX (85% O_2_) and used to quantify *CCKN1*, *SERPINE1*, *ACTA2*, *CCND1*, *ELN*, *PDGFRA*, *PDGFRB*, *TCF21*, and *TGFBR2* RNA expression by qRT-PCR. n=4; *p<0.05. F) IPA analysis indicating less pulmonary fibrosis signaling in female fibroblasts compared to male fibroblasts in HOX. The dots in heatmaps represent values that had a |Z-score| of <2 (non-significant).

In HOX, *Pdgfra* expression decreased profoundly in all fibroblasts, especially AMF (Fig. 6A and Sup. Fig. 4A). PDGFRA signaling in AMF plays a critical role in secondary septation and alveolarization^61,68^. ELN, which forms looping elastic bundles to surround alveolar ducts, and forms a continuous sheet or network in blood vessel walls ^69^, is highly expressed in AMF and VSMC in RA, and its expression in these cells dramatically decreased in HOX (Sup. Fig. 4A). ACTA2 expression in myofibroblasts is considered to be a key determinant of remodeling and disease progression in lung fibrosis^46,70^. *Acta2* expression was upregulated in DMF in HOX (Fig. 6A and Sup. Fig. 4A). *Notch3* and *Pdgfrb* were highly expressed in smooth muscle (like) cells in RA, and their expression was repressed in HOX (Fig. 6A and Sup. Fig. 4A). Notch3 signaling in mural cells is important for cell proliferation, differentiation and survival and vascular integrity^71–73^. These patterns of changes in stromal cells could contribute to the interrupted secondary septation and alveolarization, and damaged integrity of the vasculature in HOX, which are consistent with the reported increased fibrosis in human BPD ^74^.

Interestingly, a HOX-induced DIF (iDIF) subcluster emerged in HOX (Fig. 6A). Dot plot shows Haptoglobin (HP) and serum amyloid A3 (SAA3) were specific markers for this population while DIF marker genes, such as *Wnt2*, *Tcf21* and *G0S2*, were less expressed in iDIF (Fig. 6A). Volcano plot comparing iDIF to DIF in HOX shows significantly increased expression of TGFB signaling and its downstream genes (*Pdgfrb*, *Ccnd1*, *Col12a1*, *Runx1*, *Timp1*) ^75–79^, and less expression of *Inmt*, a gene enriched in DIF (Fig. 6B)^80^. DIF, as lipid storing fibroblasts, provide lipids to AT2 cells for surfactant production^81^. MLX interacting protein like (MLXIPL) signaling induces lipogenesis in adipocytes and liver, and PPARG signaling is involved in adipocyte lipid metabolism and inhibits the conversion of lipofibroblasts to myofibroblasts ^82–84^. IPA analysis revealed that both MLXIPL and PPARG upstream regulators were repressed in HOX, and even more repressed in the iDIF subcluster (Fig. 6C). Consistent with this finding, we noted upregulation of *Runx1*, a transcription factor recently described to regulate the differentiation of fibroblasts into myofibroblasts, which produce matrix proteins, thus contributing to fibrosis ^80^.

We validated scRNA-seq data in independent mouse lung samples exposed to HOX or RA [n=4/group]. qRT-PCR confirmed increased expression of *Acta2* and decreased expression of *Pdgfra*, *Pdgfrb*, *Notch3* and *Eln* in the whole lung in HOX (Fig. 6D). We wanted to determine whether changes in mouse lung fibroblasts can be recapitulated in human fibroblasts in a cell autonomous manner. We obtained human lung fibroblasts and immortalized them. We confirmed fibronectin expression in immortalized fetal human pulmonary fibroblasts (HPF-Im) using immunofluorescence staining (Sup. Fig. 4B). HPF-Im were exposed to 85% oxygen (HOX) or normoxia for seven days, and the cells were used to perform qRT-PCR which demonstrated that HOX repressed the expression of *PDGFRA*, *PDGFRB* and *ELN*, while upregulating *TCF21* expression (Fig. 6E). TP53 target genes (*SERPINE1* and *CDKN1A*) TGFβ signaling marker genes (*TGFBR2, SERPINE1* and *CDKN1A*) and fibrosis signaling marker genes (*CCND1* and *SERPINE1*) were upregulated in HOX, but their shared downstream gene *ACTA2* didn’t change (Fig. 6E). Our HPF-Im data is mostly consistent with our mouse lung data in HOX. The increase in *TCF21* with decrease in *ACTA2* in HPF-Im with HOX is likely representative of a shift towards a *Tcf21*^+^DIF signature, which is consistent with an increase in this population with HOX in the mouse lung (Fig. 6A).

Next, we compared signaling pathways between female and male mouse lung fibroblasts after HOX treatment using the gene expression data from scRNA-seq. IPA canonical pathway analysis showed less pulmonary fibrosis signaling pathway in four types of fibroblasts in female in HOX (Fig. 6F). Less pulmonary fibrosis signaling in female fibroblasts may lead to reduced lung fibrosis in female in HOX, though this will have to be experimentally demonstrated.

### Immune cells show sex-specific changes in their transcriptome following hyperoxia

Based on *Ptprc* and individual marker genes, 12 major clusters of *Ptprc*^+^ immune cells were identified (Fig. 7A): *Cd79a*^+^ B cells, *Cd79a*^+^/*Mki67*^+^ cycling B (cB) cells, *Cd3g*^+^ T cells, *Cd3g*^+^/*Mki67*^+^ cycling T (cT) cells, *Ear1*^+^ alveolar macrophages (AM), *Ear1*^+^/*Mki67*^+^ cycling AM (cAM), *Mafb^high^* interstitial macrophages (IM), *Ly6c2^high^* monocytes, *Xcr1*^+^ dendritic cells (DC), *Xcr1*^+^/*Mki67*^+^ cycling dendritic cell (cDC), *S100a9*^+^ neutrophils and *Cd200r3*^+^ basophils (Fig. 7B). We used IPA canonical pathways analysis to identify major transcriptomic changes filtered by the different cell types (Fig. 7C). It revealed less B cell receptor signaling, PI3K signaling, B cells activating factor signaling, CD27 signaling and leukocyte extravasation signaling in B cells. It also showed less 4-1BB signaling in T cells. Interestingly, IPA analysis demonstrated more leukocyte extravasation signaling, production of nitric oxide (NO) and reactive oxygen species (ROS), Fcy receptor-mediated phagocytosis and MSP-RON signaling in AM in HOX (Fig. 7C).

**Figure 7.**
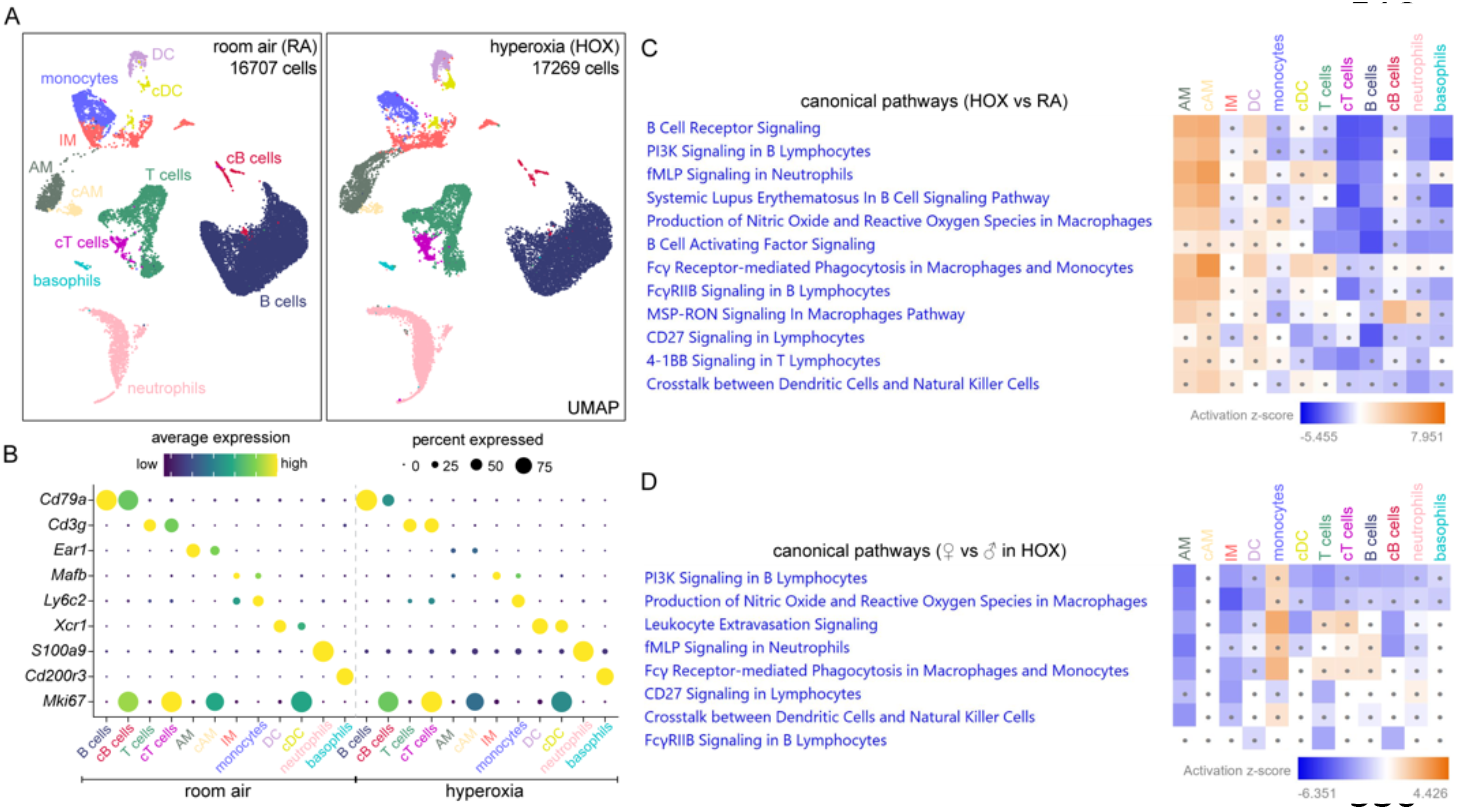
Hyperoxia induces sex-dependent changes in immune cells. A) UMAPs of immune cells showing distinct clusters of B cells, cycling (c)B cells, T cells, cTcells, AM, cAM, IM, monocytes, DC, cDC, neutrophils, and basophils in RA and HOX. B) Dot plot showing marker genes used to identify each immune cell cluster. C) IPA analysis revealing less B cell receptor signaling, B cells activating factor signaling, CD27 signaling and leukocyte extravasation signaling in B cells, but upregulated production of NO and ROS, Fcy receptor-mediated phagocytosis, and MSP-RON signaling in AM in HOX. D) IPA analysis indicating less upregulated leukocyte extravasation signaling, production of NO and ROS and Fcy receptor-mediated phagocytosis in female AM compared to male AM in HOX. (C-D) Dots in heatmaps represent values that had a |Z-score| of <2 (non-significant). AM, alveolar macrophages; IM, interstitial macrophages; DC, dendritic cells.

Next, we compared canonical pathways between female and male in HOX. IPA analysis revealed less PI3K signaling in B cells and less CD27 signaling in T cells. Remarkably, it also revealed female AM had less upregulation of leukocyte extravasation signaling, production of NO and ROS, and Fcy receptor-mediated phagocytosis in HOX (Fig. 7D). NO and ROS produced by activated macrophages protect hosts against a broad spectrum of pathogens and stimulation^85,86^, while Fcy receptor-mediated phagocytosis plays a key role in clearing pathogens^87^. AM are the first line of defense in the lung against airborne pathogens, clearing injured cells and debris ^88,89^, but they also contribute to pathogenesis and pulmonary fibrosis by releasing excessive cytokines, oxygen radicals and proteases^6,90–92^. Our IPA analysis revealed less AM activation and phagocytosis in females compared to males in HOX.

## DISCUSSION

To investigate how hyperoxia programs transcriptional networks regulating cellular identity plasticity, injury and repair in the neonatal lung, and the impact of sex on transcriptomic signatures, we combined scRNA-seq with validation studies in independent mouse lung samples and neonatal human lung cell lines. HOX acutely induces AT1 injury, and it is postulated that AT2 cells serve as the progenitor cells that repopulate AT1 cells upon injury ^93^. In the developing mouse lung, we identified Lyz1^+^ AT2, AT2, cAT2 cells and another AT2 cluster, which we labelled mAT2. The Lyz1^+^ AT2 cluster had previously been noted by Hurskainen et al. ^7^ in their study describing cellular populations after HOX. However, mAT2 cells have not been described in the neonatal lung before. This cell population showed enrichment of mitochondrial biogenesis genes such as *mt-ND1* and *mtND2*, while having decreased expression of *Sftpc*, *Etv5* and *Abca3*. In HOX, we first noted and then confirmed by IF that AT2 cells assumed an ‘inflamed’ phenotype with increased *Lcn2* expression, an observation Hurskainen et al. also noted in their study. The striking finding among AT2, was the emergence of DATP population (*Sftpc^low^*/*Krt8*^+^/*Cldn4*^+^) in HOX, which was not present in RA. This population has been described as a transient stem cell progenitor state induced by bleomycin in the adult lung ^5^ but it had not been described before in hyperoxia models. IPA analysis revealed activation of IL1β and HIF1α pathway signaling in HOX-exposed AT2 cells, as reported by Choi and Nold, et al., which revealed that these pathways program AT2 towards an AT1 phenotype ^28,94^. The switch from AT2 to inflamed AT2 to DATP in the neonatal lung with HOX, suggests that AT2 have similar adaptive/regenerative mechanisms to various noxious stimuli. In primary neonatal human alveolar epithelial cells *in vitro* we showed that HOX cell-autonomously induces expression of the inflamed AT2 and DATP signatures, suggesting a conserved response across species. Scafa et al. used a brief exposure to HOX (P1-P3) to study AT1 and AT2 populations using scRNA-seq, and while they identified several AT clusters, they did not comment on the populations we identified ^95^. Whether this response is able to reconstitute the AT1 pool, and the key signaling mechanisms mediating it, are topics for future research.

In AT1 cells, we noted a distinct cluster emerge in HOX marked by increased mitochondrial biogenesis genes, which we labeled as iAT1. These iAT1 cells had decreased expression of the traditional AT1 markers and AT1-derived growth factors. We also confirmed their emergence in vivo by co-localization of mt-ND1 and PDPN in the mouse lung after HOX. Whether these cells represent an injured state of AT1 or an immature population of newly formed AT1 cells, and the specific mechanisms governing their emergence - whether from DAPT, AT2/AT1 or from AT2 cells - needs to be further investigated. Comparing alveolar epithelial cells between females and males in HOX to determine transcriptomic changes regulating resilience of female pups to hyperoxia suggested less EMT, pulmonary fibrosis and necroptosis signaling in females.

We identified two populations of capillary EC as reported by other groups ^3,30–32^. Cap2 appear to be specialized for gas exchange but do not proliferate, whereas Cap1 function as a progenitor cell for Cap2 during lung development and injury recovery^32,33^. In our study, *Car4*^+^ Cap2 and Car4 expression in Cap1 increased after HOX, indicating a reparative capillary response. Hurskainen et al., also found a significant increase in the Cap2 population in HOX^7^, which is consistent with our data, but they did not address sex-specific endothelial signatures. We also noted the emergence of iCap2, a new subpopulation of Cap2 cells not described before. These cells expressed *Car4* and *Apln* to a lesser extent than bona fide Cap2, but not *Plvap* or *Aplnr*, the Cap1 markers. Other distinguishing features of this population were decreased *Kdr* and *Nrp1*, which are key mediators of VEGF signaling ^96^. Whether iCap2 cells are injured or newly formed Cap2, remains to be deciphered, but decrease in lumenization marker *Icam2* would suggest they represent an immature state. ScRNA-seq showed more *Car4* expression in female Cap1 than male Cap1 cells in HOX. Validation studies further confirmed higher *Car4* mRNA and protein expression in female neonatal mice. We did not have the tools to isolate Cap2 from Cap1 to validate increased CAR4 in the Cap1 population, but this sex-dependent difference was so pronounced that we could detect it in whole lung lysates. Altogether, these data might indicate a more robust EC regenerative response in females ^33^. Expression of both Cap2 and Cap1 markers, such as *APLN*, *CA4*, *APLNR* and *PLVAP*, along with *KDR* was repressed by HOX in primary canalicular stage HPMEC, whereas the addition of human fetal lung fibroblasts in a co-culture system upregulated gene expression of both Cap1 and Cap2 markers. These data suggests that the EC recovery from HOX requires paracrine input from fibroblasts and AT1. We speculate that AT1-derived VEGFA might be required for selective Cap2 expansion in HOX, as it is a known key factor for emergence of *Car4*^+^ Cap2 during development^31^. Moreover, although scRNA-seq shows an increase in *Car4*^+^ Cap2 cells, IF shows a decrease in CAR4 staining. This could be explained by the formation of new Cap2 cells in HOX, which would potentially have shorter extensions as they start to develop, visually leading to a less dense CAR4 network. Both Cap2 EC and AM express *Car4* under normal conditions. However, we believe that the main source of increased *Car4* expression in mouse whole lung experiments in HOX comes from expansion of Cap2 vessels, since *Car4* seems unchanged in scRNA-seq data comparing AM between both conditions.

We defined stromal mesenchymal cells as previously described in a recent paper^36^. Among these subpopulations, *Pdgfra^+^/Wnt2*^+^/*Tcf-21^high^/G0s2^high^* DIF, *Pdgfra^high^/Lgr6^-^/Hhip*^+^/*Cdh4*^+^ AMF and *Pdgfra*^+^/*Twist2*^+^/*Il33*^+^/*Col1a1^high^*/ *Col1a2^high^* PIF were similarly defined as *Col13a1*^+^ fibroblasts, myofibroblasts and *Col14a1*^+^ fibroblasts, respectively, in another neonatal HOX study^7^. SAA3 was enriched in interstitial fibroblasts in HOX, which is consistent with Hurskainen study^7^. *Pdgfra* expression was repressed in all fibroblasts in HOX, which is also consistent with scRNA-seq data from Hurskainen et al^7^. We also identified a new subcluster of DIF in HOX, iDIF, in which expression of DIF marker genes was reduced, but TGFB downstream genes were upregulated. Incidentally, upregulation of *Runx1* and suppression of *Pparg* in iDiF also suggest these cells might differentiate into collagen-synthesizing myofibroblasts, and contribute to the fibrotic response seen in HOX and BPD^97^. EC induce differentiation and maturation of mural cells via NOTCH3^98,99^, whose expression was decreased in mural cells in HOX. PDGFRA signaling in AMF plays a critical role in alveolar septation, and *Pdgfra^-/-^* mice exhibit alveolar simplification^61^. Repressed expression of *Pdgfra* and *Eln* in AMF could be a key cause of disrupted alveolar development in HOX. We revealed that *Pdgfra* and *Eln* mRNA expression was repressed in HOX, while TGFBR2 and SMAD3 protein expression and phosphorylation was upregulated in the whole lung. We further confirmed this in neonatal human fibroblasts *in vitro*. HOX repressed *PDGFRA* and *ELN* expression, which are critical to secondary septation, whereas HOX upregulated the expression of *TGFBR2* and *ACTA2*, key determinants of remodeling and disease progression in lung fibrosis^46,70^. IPA analysis showed less pulmonary fibrosis signaling in fibroblast clusters in female mice exposed to HOX, which might explain decreased BPD vulnerability in female infants.

While our focus was on epithelial, endothelial and mesenchymal cells that directly contribute to alveolarization, we also examined the immune cell population. Our results did not identify any new populations in HOX but IPA analysis revealed HOX stimulated important pro-inflammatory signaling pathways in AM. The expansion of activated AM population in HOX could represent monocyte-derived alternatively activated macrophages^100^. Interestingly, HOX-induced leukocyte extravasation signaling, production of NO and ROS, and Fcy receptor-mediated phagocytosis was decreased in females compared to males, which may be relevant to decreased BPD incidence in female infants, as AM activation is known to interrupt lung development^6^.

In conclusion, our study shows that hyperoxia programs distinct cellular and transcriptomic changes that signal injury and reparative responses in the neonatal lung. The emergence of DATP and iCap2 most likely represent reparative responses, while whether iAT1 and iDIF populations represent injury states or newly generated immature populations remain to be discerned. Female mice showed signatures of decreased fibrosis signaling, a more robust EC iCap2 response and decreased macrophage activation, all of which could contribute to decreased vulnerability of females to HOX injury. Interestingly, neonatal human alveolar epithelial cells and fibroblasts exhibit cell-autonomous reparative gene expression signatures in HOX that resemble our mouse data, while cell-autonomous EC responses mimic an injury response, which is rescued with fibroblast co-culture. While we report several novel and interesting findings, a lack of insight into the transcriptional mechanisms that program these distinct cell-type specific changes remain a limitation. However, our findings lay a foundation for future mechanistic studies that can target the cell types and signaling pathways here identified.

## MATERIAL AND METHODS

### Study Approvals

Lab experiments were reviewed and approved under the University of Missouri-Kansas City IBC, protocol number 18-28 and the Children’s Mercy Research Institute IBC, protocol number 025. Animal experiments were reviewed and approved under the University of Missouri-Kansas City IACUC, protocol number 1510-03.

### Mice (*Mus musculus*)

Mice were housed and treated with hyperoxia at the University of Missouri-Kansas City animal facility. Mice were kept in a 12-hour light/dark schedule as per the facility animal housing protocol described before ^38,67^. *WT* C57BL/6 (Charles River) mice were used in the study. For hyperoxia, pups with dam were placed in cages in a hyperoxia chamber for 14 days from P1 to P14. We used 3 females and 3 males/experimental condition for scRNA-seq, and 4 females and 4 males for validation studies. The enclosure was flushed with oxygen and the concentration was continuously monitored and controlled with a Pro-ox oxygen controller (BioSpherix) to maintain 85% O_2_ throughout the study period. Control mice were housed similarly but were not exposed to hyperoxia. Two dams were switched daily between hyperoxia and room air. Pups were euthanized using a 100mg/kg intraperitoneal injection of pentobarbital, exsanguinated after cessation of heartbeat, and the lungs were harvested as described below.

### Cell dissociation

Lungs from experimental C57BL/6 mice were inflated with an enzyme mix consisting of the following: Leibovitz media (Invitrogen) 2mg/mL of collagenase type 1 and 2mg/mL of elastase (Worthington Biochemical). Lungs were then chopped with a razor blade into fine chunks, and placed in a prewarmed digestion enzyme mix - inflation enzyme mix plus 0.5mg/mL DNase 1 (Worthington Biochemical). They were placed in an incubator at 37C and rotated for 15min; next they were mechanically digested with a 1ml pipette tip. The digestion mix was placed back into the incubator and rotated for another 15min, after which the digestion was stopped with 20% FBS and the lysate was filtered through a 70um filter (Fisher Scientific). The lysate was then centrifuged for X min at X rpm at X temperature, and incubated with 1X red cell lysis buffer (Sigma-Aldrich) for 3 minutes. The sample was centrifuged for X at X, and the supernatant was discarded; the sample was then resuspended in Leibovitz media containing 3% FBS, and filtered as before. Dissociated cells were counted after centrifugation and used for scRNA-seq.

### scRNA-seq

12 C57BL/6 pups were euthanized using 100 mg/kg intraperitoneal injection of pentobarbital at P14. Fresh lung single cell digests furnished > 10-20×10^6^ cells/mouse with >80% viability.12,000 cells were loaded into a 10X Genomics Chromium Single Cell A Chip (10X Genomics; PN-120236), and scRNA-seq was performed using Chromium Single Cell 3’ Library & Gel Bead Kit v2 (10X Genomics; PN-120237) following the manufacturer’s protocol ^67^. Briefly, single-cell gel beads-in-emulsion (GEMs) were produced by running a loaded Chromium Single Cell A Chip on a Chromium Controller instrument (10X Genomics). After GEM-reverse transcription (GEM-RT), GEMs were collected, and cDNA was amplified and cleaned up using SPRISelect (Beckman Coulter; B23318). Indexed libraries were prepared using the Chromium Single Cell 3’ Library & Gel Bead Kit v2 (10X Genomics; PN-120237) and sequenced on a NovaSeq 6000 (Illumina) to obtain a sequencing depth of 1.50E+08 paired-end reads per library. Raw sequencing data was processed through bcl2fastq2 (Illumina), resulting in 1 fastq file per library. Individual 10x libraries were processed through Cell Ranger v3 0.2, and resulting count matrices imported into Seurat v4 1.1 (https://satijalab.org/seurat). Technical variability was reduced using Seurat’s ‘sctransform’, and quality control metrics were analyzed per sample. The data was then normalized, scaled, identification of variable genes and PCA were performed, followed by dimensionality reduction. Clusters were analyzed with FindAllMarkers function in order to identify marker genes for each cell lineage. Each lineage was then subsetted into a new object and reanalyzed. Finally, volcano plots were generated using the library EnhancedVolcano with a Log2 fold change cut off of 1, and a p value cut off at 10e-6^101^. Raw data have been deposited in GEO under the accession number X (pending).

### IPA analysis

Gene expression data of scRNA-seq were obtained by differentially expressed genes were defined as p<0.05 as assessed using unpaired-samples t-test. The IPA system (version 73620684, Ingenuity Systems; Qiagen China Co., Ltd.) was used for subsequent bioinformatics analysis, which included canonical pathways, upstream analysis, diseases and functions, and regulator effects. For analyses, the −log (P-value) >1.3 [p<0.05] was taken as threshold and a Z-score of ± 2 was defined as the threshold of significant activation/repression.

### Cell culture and immortalized embryonic human lung fibroblasts (HPF) generation

Immortalized embryonic human pulmonary microvascular endothelial cells (HPMEC-Im) was generated and grown as done in previous studies ^67,102^. Primary fetal human pulmonary fibroblasts (HPF) and primary fetal pulmonary alveolar epithelial cells (HPAEpiC) were purchased from ScienCell (Carlsbad, CA). We followed the company’s protocol to culture the cells. Immortalized HPF were generated using Lentivirus containing SV40 large T antigens (ABM, Richmond, BC, Canada). The Immortalized HPF (HPF-Im) was verified with fibronectin staining (Sup Fig. 3). Cells were cultured in a sealed chamber containing 85% O_2_ and 5% CO_2_ for 2-day (for AT cells) or 7-day (HPAEpiC and HPF-Im) hyperoxia treatment.

### Quantitative reverse transcription PCR (qRT-PCR)

Total RNA was extracted from mouse lung or cultured cells using the PureLink RNA Mini Kit (Invitrogen) and cDNA was synthesized from 1μg of RNA using an iScript cDNA synthesis kit (Bio-Rad), according to manufacturer’s instructions. qRT-PCR was run on a ViiA 7 with SYBR green mastermix (Biotium). 18S was used as the human cell housekeeping gene and *Ywhaz* was used as the mouse housekeeping gene. The relative gene expression was calculated using the Pflaffl method.

### Immunoblotting

Immunoblotting for quantifying changes in protein expression was performed as previously described^67^. In brief, mouse lung tissue was homogenized in RIPA lysis buffer containing commercially available protease and phosphatase inhibitors (Sigma) after RA and HOX treatment, with the clarified lysates used for western blotting (WB). Densitometry was performed using ImageJ Software (NIH) and changes were normalized to YWHAZ. Primary antibodies are described in supplementary table 2.

### Immunostaining

Immunostaining of mouse lung and cultured cells was done as in our previous study ^67^. Briefly, the lungs of the mouse pups were fixed in 4% formaldehyde and embedded in paraffin. Cultured cells were grown on coverslips and then fixed with 4% formaldehyde. Cells or deparaffined lung sections were incubated with first antibodies after blocking at 4 °C overnight, and then incubated with corresponding second antibodies at 25 °C for 1 hour. Several washing with 0. 1% tween in PBS after incubating first and second antibody and then sealed the slides with Vibrance^®^ Antifade Mounting Medium with DAPI. Images were taken using ZEISS LSM 510 confocal microscope.

### Statistical Analysis

The statistical analysis was performed as discussed in our previous study ^67^. Briefly, data are presented as mean ± SD or median with interquartile range. P < 0.05 was considered significant. For cell culture experiments, data are from a minimum of three independent experiments with adequate technical replicates used for quantification. All animal data were obtained in littermate controls. For animal experiments, a minimum of four animals were used for each experimental group with adequate technical replicates used for quantification. For all data, we initially examined whether distribution of data was Gaussian using the D’Agostino-Pearson omnibus normality test. If data were normally distributed, then ANOVA with a post-hoc Tukey test was used for analysis. If data did not meet Gaussian assumptions, a Mann-Whitney U-test was used for analysis. Comparisons between two groups were made by a 1-sample, 2-tailed Student t-test for parametric or nonparametric data. For most analysis, fold-changes were calculated related to expression/changes in untreated controls. Statistical analysis and graphs were generated using Graphpad Prism 9.0.

## Acknowledgments

HM, SX, SM and VS were supported by 1R01HL128374-01 (VS). LVE was supported by 5K99HL155845; JC was supported by R01HL153511 and R01HL130129.The authors thank Aparna Venkataraman from Children Mercy Hospital, Kansas City for critical input and Wei Yu, PhD from Children Mercy Hospital, Kansas City for producing immortalized cell lines. We also thank Qiagen for their permission to use images generated with their IPA software.

