## Supplemental material for "Neonatal hyperoxia induces sex-dependent pulmonary cellular and transcriptomic changes in an experimental mouse model of bronchopulmonary dysplasia"

### Sup. Fig. 1 Lung images and quantification

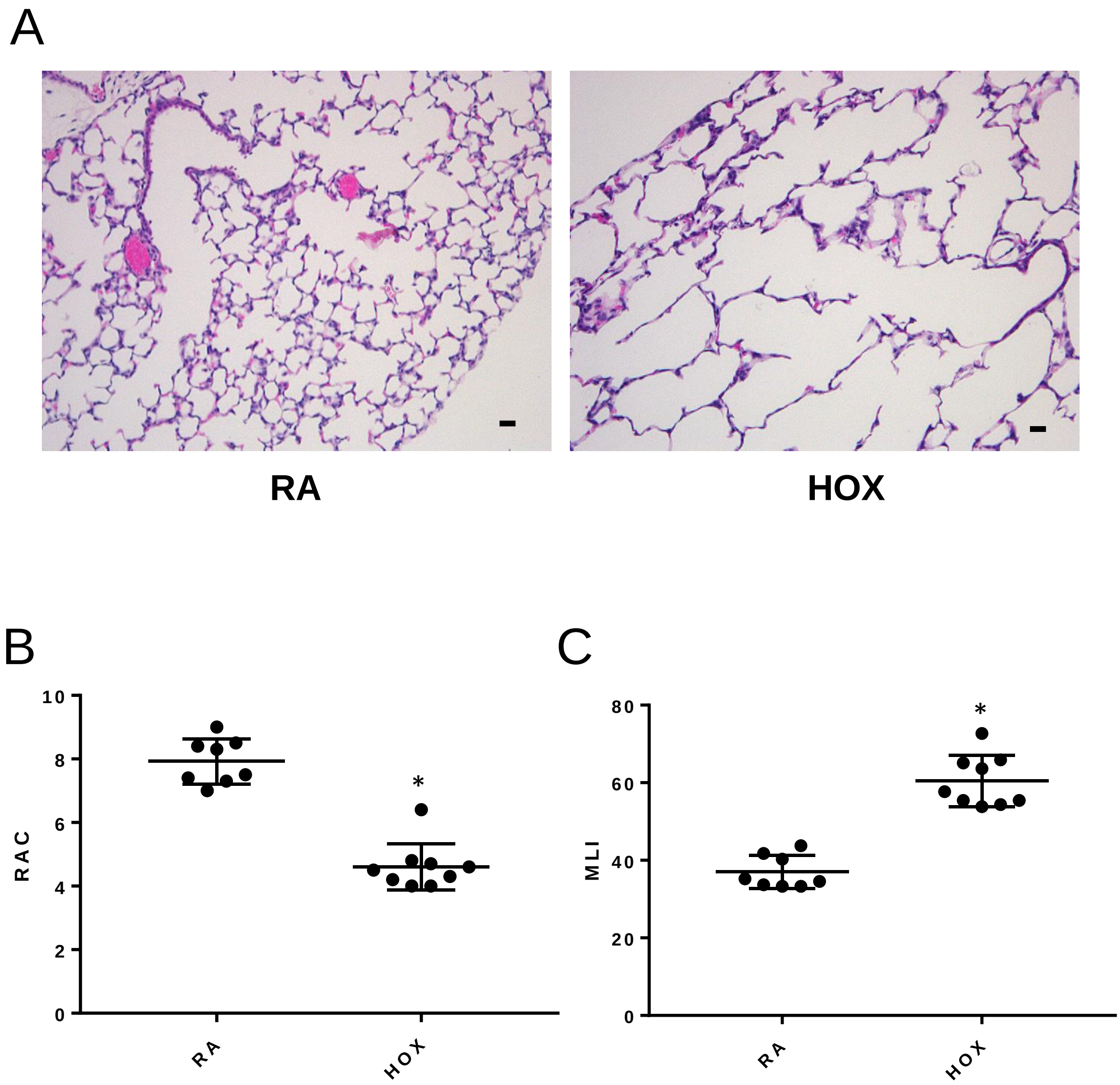

### Sup. Fig. 2 AT diversity and alteration in hyperoxia

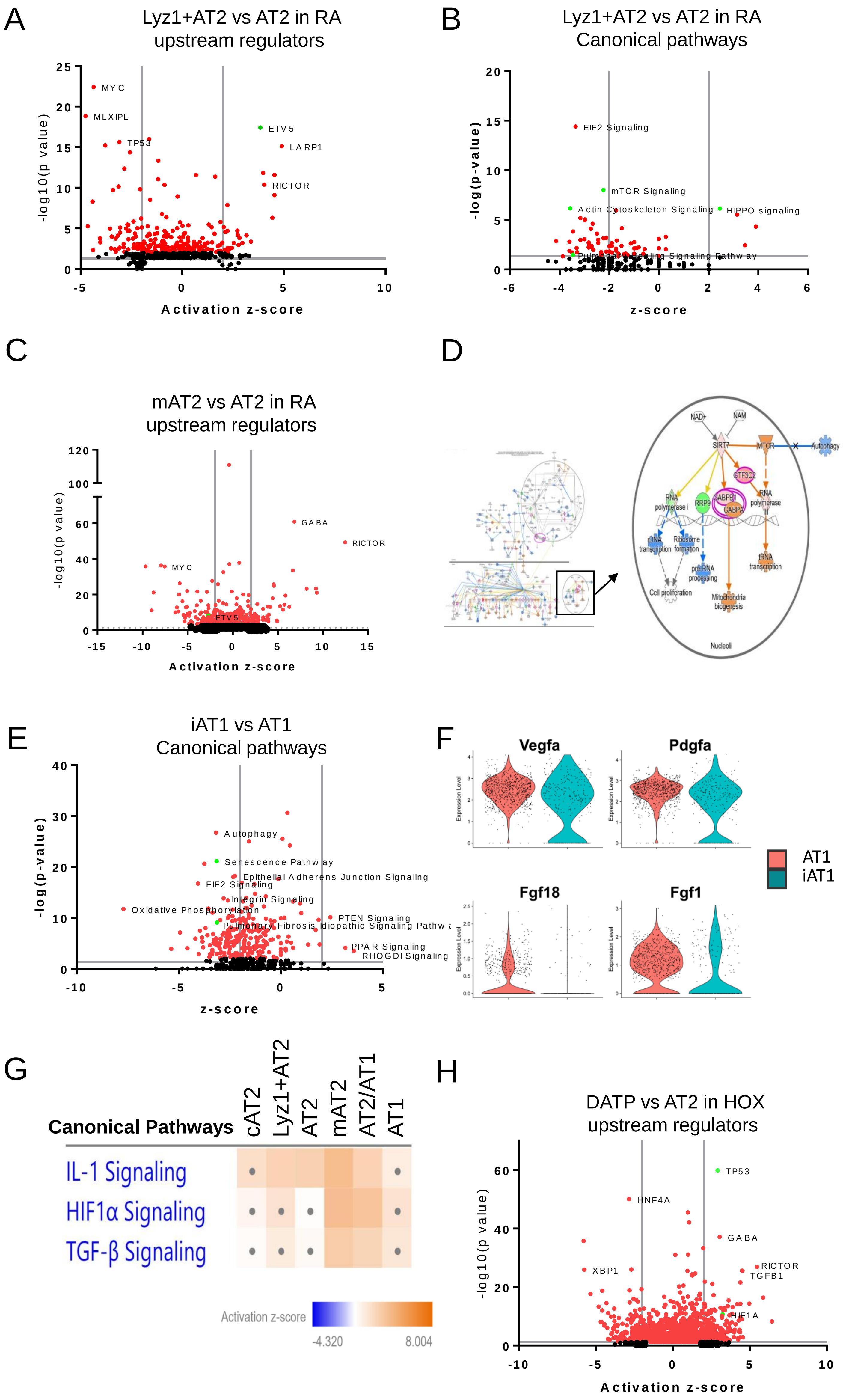

### Sup. Fig. 3 EC diversity and alteration in hyperoxia

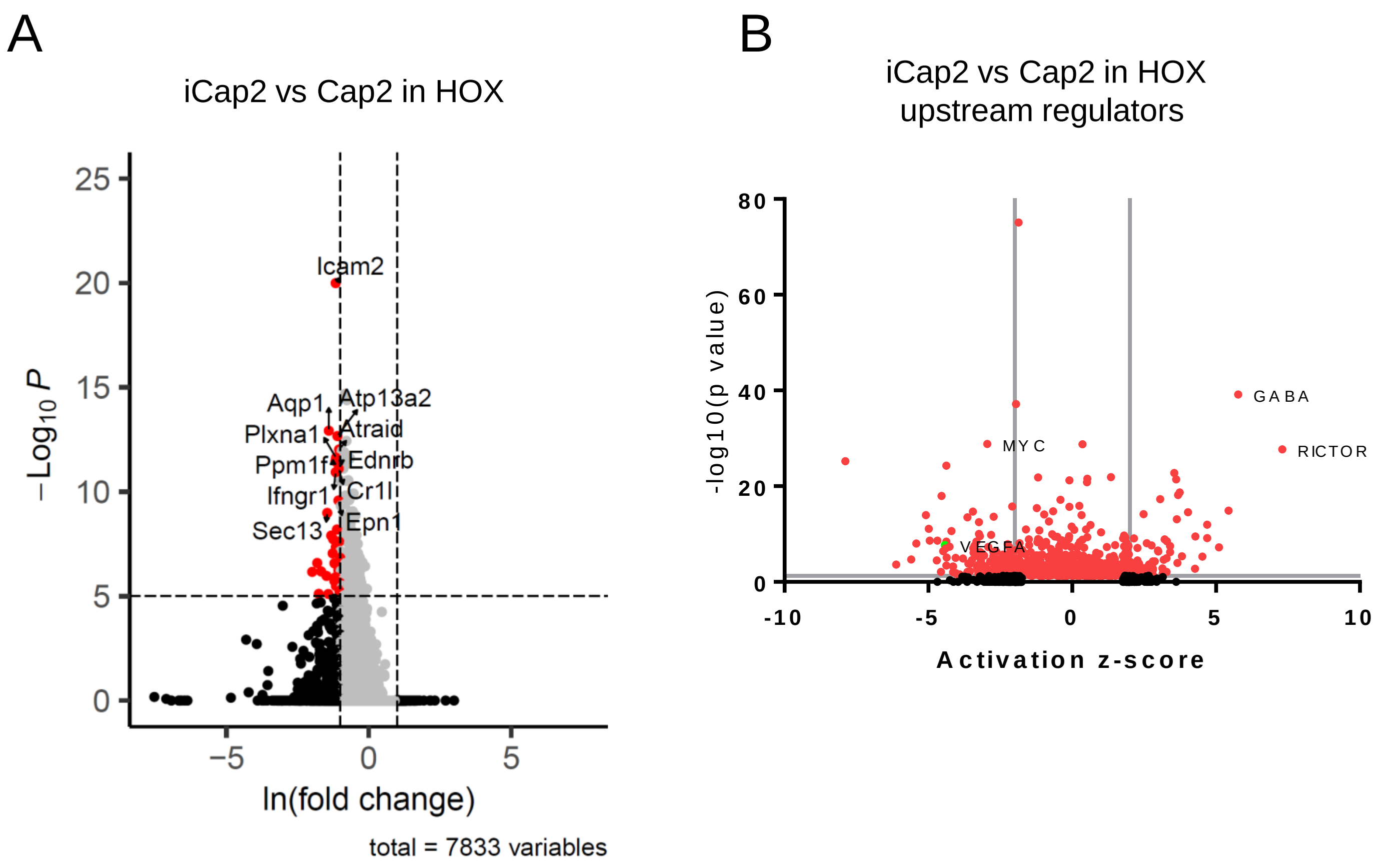

### Sup. Fig. 4 Stromal cell alteration in hyperoxia

A

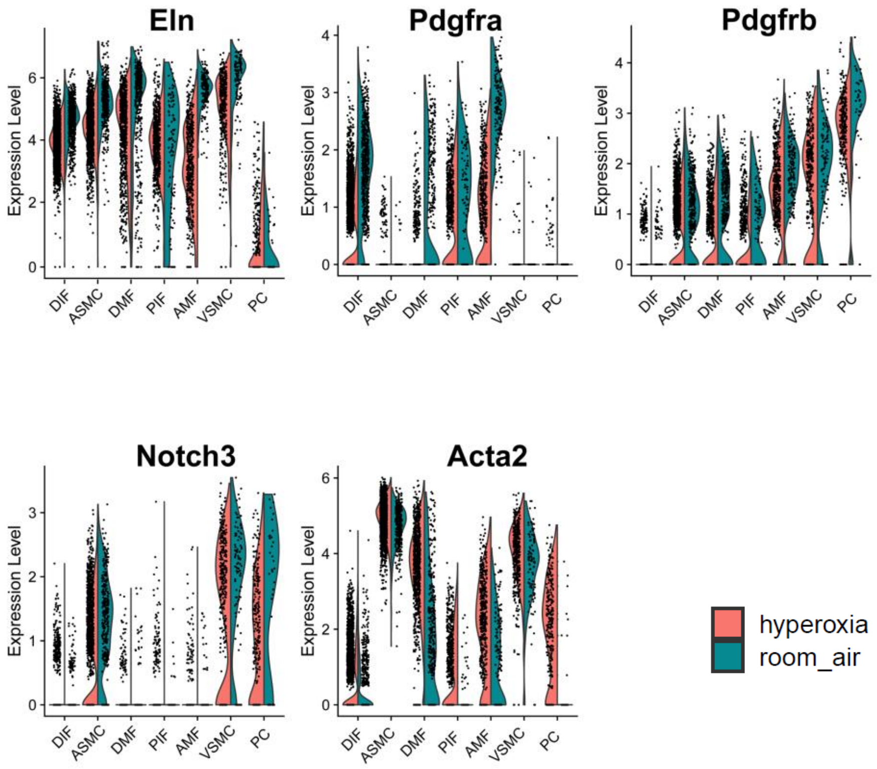


B

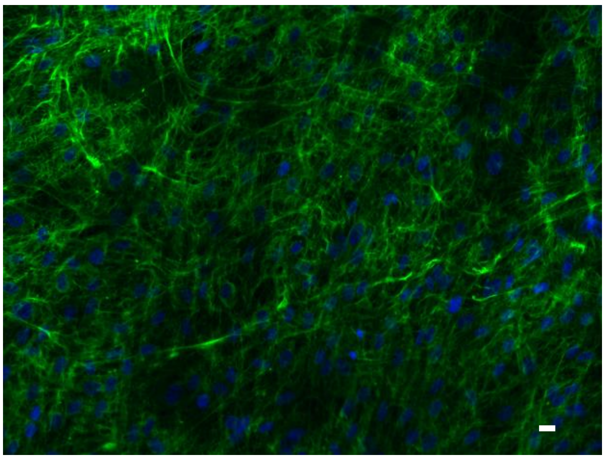

Fibronectin/DAPI

**Supplementary Figure Legends**

**Sup. Fig. 1: Hyperoxia causes alveolar simplification in mouse lungs**. Neonatal mouse pups were exposed to 85% oxygen (HOX) or room air (RA) from P1-P4. Inflation-fixed lungs were used for measuring radial alveolar counts and mean linear intercepts. (A-C) Images of H&E-stained lungs harvested from HOX or RA-treated P14 pups (A), with graphic representations of radial alveolar counts (RAC) (B) and mean linear intercepts (MLI) (C) shown. Scale bar is 25μm, n≥6 per group, *p<0.01.

**Sup. Fig. 2: Alveolar epithelial cell diversity and transcriptomic changes in hyperoxia. (**A-C) Volcano plots showing IPA comparisons of upstream regulators between Lyz1$^{+}$ AT2 vs AT2 cells in RA (A), canonical pathways between *Lyz1*$^{+}$ AT2 vs AT2 in RA (B) and upstream regulators between mAT2 vs AT2 in RA (C). Significantly up/downregulated pathways are labeled in red, and some biologically relevant pathways and regulators have been highlighted in green. D) Diagram showing IPA canonical pathway analysis for mitochondria biogenesis (right) and sirtuin signaling (left) in mAT2 compared to AT2 in RA (downloaded from IPA with permission). Pink/orange indicates upregulation while green/blue indicates downregulation. E) Volcano plot of IPA canonical pathway analysis between iAT1 vs AT1 cells in HOX. Significant changes are shown in red, and some relevant changes are in green. F) Violin plots showing decreased expression of *Vegfa*, *Pdgfra*, *Fgf1,* and *Fgf18* in iAT1 vs AT1 in HOX. G) Heatmap comparing IPA upstream analysis in *Lyz1*+ AT2, AT2, mAT2 and AT2/AT1 clusters in HOX vs RA. H) Volcano plot of IPA upstream analysis between DATP vs AT2 in HOX. Significant changes are marked with red or green colors. │Z-score│ cutoff=2 (<2 non-significant) for all volcano plots.

**Sup. Fig. 3: Endothelial cell diversity and transcriptomic changes in hyperoxia.** A) Volcano plot of iCap2 vs iCap2 in HOX showing differential gene expression patterns in iCap2. B) Volcano plot of IPA comparing iCap2 vs Cap2 in HOX. Significantly up/downregulated upstream regulators are labeled in red, and some biologically relevant pathways have been highlighted. │Z-score│ cutoff =2 (<2 non-significant).

**Sup. Fig. 4: Stromal mesenchymal cell in hyperoxia.** A) Violin plots depicting changes in gene expression of *Eln*, *Pdgfra*, *Pdgfrb,* and *Acta2* in distinct populations of stromal cells in RA (blue) vs HOX (red). B) Fibronectin (green)/DAPI (blue) immunofluorescence in immortalized human primary fibroblasts (HPF-Im). Scale bar = 10μm.

**Supplementary Table**

| Anti- | Host | Company | Catalogue No. |
|---|---|---|---|
| CAR4 | Goat | ThermoFisher Scientific | PA5-47312 |
| PLVAP | Rat | BD Biosciences | 553849 |
| ERG | Rabbit | Abcam | ab92513 |
| SPC | Rabbit | Abcam | ab90716 |
| PDPN | Rat | Abcam | ab256559 |
| KRT8 | Rat | DSHB | AB_531826 |
| LCN2 | Rat | Novus Biologicals | NBP1-05183 |
| MKI67 | Rat | ThermoFisher Scientific | 11-5698-82 |
| Fibronectin | Rabbit | Proteintech | 15613-1-AP |
| SAA3 | Rat | Abcam | ab231680 |
| Mt-Nd1 | Rabbit | ThermoFisher Scientific | PA5-120599 |
| CAR4 (WB) | Rabbit | ThermoFisher Scientific | PA5-86930 |
| PLVAP (WB) | Rabbit | ThermoFisher Scientific | PA5-115774 |
| TGFBR2 | Mouse | ThermoFisher Scientific | 66636-1-IG |
| pTGFBR2 | Rabbit | ThermoFisher Scientific | PA5-105920 |
| SMAD3 | Rabbit | ThermoFisher Scientific | 51-1500 |
| pSMAD3 | Rabbit | ThermoFisher Scientific | 600-401-919 |
| YWHAZ | Rabbit | ThermoFisher Scientific | PA5-80245 |
